# Topological analysis reveals state transitions in human gut and marine bacterial communities

**DOI:** 10.1101/584201

**Authors:** William K. Chang, Dave VanInsberghe, Libusha Kelly

## Abstract

Microbiome dynamics influence the health and functioning of human physiology and the environment and are driven in part by interactions between large numbers of microbial taxa, making large-scale prediction and modeling a challenge. Here, using topological data analysis, we identify states and dynamical features relevant to macroscopic processes.We show that gut disease processes and marine geochemical events are associated with transitions between community states, defined as topological features of the data density. We find a reproducible two-state succession during recovery from cholera in the gut microbiomes of multiple patients, evidence of dynamic stability in the gut microbiome of a healthy human after experiencing diarrhea during travel, and periodic state transitions in a marine *Prochlorococcus* community driven by water column cycling. Our approach bridges small-scale fluctuations in microbiome composition and large-scale changes in phenotype without details of underlying mechanisms, and provides a novel assessment of microbiome stability and its relation to human and environmental health.

## Introduction

Complex microbial ecosystems (‘microbiomes’) inhabit a diversity of environments in the biosphere, including the global ocean [47], soil [13], and the human gut [48]. Large-scale alterations in the composition of microbiomes is often associated, whether as driver or consequence, with environmental processes such as seasonal geological cycling and nutrient fluctuations [15]; physiological processes such as menstrual cycles [16]; and clinical phenotypes such as irritable bowel syndrome [2]. Analysis and prediction of the large-scale dynamics of microbiome composition is thus a pressing issue in multiple fields of study.

As with many biological systems, understanding of the dynamics of microbiomes is complicated by their high dimensionality. Numerous variables define the state of a microbiome; these include frequencies of microbial taxa and their genetic alleles, which are decoupled due to genomic plasticity and horizontal gene transfer [36, 38], and environmental conditions such as temperature, pH, and biochemical concentrations. A microbiome thus has a vast number of potential configurations in which it may, in principle, fluctuate on a short time scale. By contrast, systemic phenotypes, such as human gut infections or aquatic algal blooms, persist for much longer than bacterial generation time, and community compositions may be diverse within a phenotype [15]. Furthermore, due to the diverse biology of microbiomes across habitats, it may be desirable to have a quantitative framework that can be generalized across biological systems.

One approach to analyzing microbiome dynamics has been to infer the network of underlying pairwise interactions between taxa by calculating the inverse covariance matrix from time series data, often as a basis for modeling population dynamics using Lotka-Volterra equations [14, 28, 46]. Such approaches are useful for predicting fine-grained taxon-taxon interactions of importance, and are challenged by the compositional nature of microbiome data [44] and possible role of higher-order interactions [3]. Notably, it is impossible to fit Lotka-Volterra models to compositional data without information regarding the total population size [26]. A complementary coarse-grained approach is to cluster samples according to compositional similarity, and conceptualize dynamics as stochastic transitions between clusters [1, 9]. Such approaches can be used to identify large-scale shifts in compositional state, with the implicit assumption that each temporal sample can be assigned to one of a finite number of discrete categories.

In our approach to microbiome dynamics, we were motivated by the concept of potential landscapes in physics. The potential landscape formalism considers a high-dimensional phase space, in which coordinates represent system states, and system dynamics correspond to trajectories through phase space. The dynamics are envisioned as being influenced by features of a landscape in phase space, the height of which corresponds to the value of a potential energy function: for example, local minima of the potential may represent stable states, and valleys probable dynamics of the system. In biology, the potential landscape and related concepts have proved useful in theoretical and experimental studies of ecological dynamics [6, 7, 41]; cell phenotypes in differentiating stem cells [50, 52] and cancer cells [25, 30]; and states of brain activity [20].

In principle, potential landscapes predict an inverse relationship between the value of the potential and the probability of observing the corresponding system state, and thus between the potential in a region of phase space and the density of observations in that region. In reality, certain landscape features and dynamics may lead to the persistence of transient states and the illusion of stability [22, 35], and strong external perturbations may cause the dynamics to deviate from those predicted by the potential landscape, in particular in biological applications. For example, perturbations to the gene expression of a differentiating stem cell may cause it to lose or fail to attain a differentiated phenotype [24]. While the potential landscape formalism may not be directly applicable to microbiomes due to the open nature of the system and rapid turnover relative to currently-practical sampling frequency, we speculated that creating a representation of the density of data points in the compositional phase space of microbial ecosystems could lead to useful insights for analyzing, and eventually predicting, microbiome dynamics. Specifically, we hypothesized that local maxima of the data density could form a basis by which to infer characteristic metastable states of microbiome composition, allowing the association of observations with states and the representation of dynamics as metastable state transitions while retaining the continuity of the underlying phase space.

To characterize features of the microbial phase space, we used topological data analysis (TDA), specifically the Mapper algorithm [39, 45], which has recently found application in microbiome research [31]. TDA is a class of methods for inferring properties of data, represented as a point cloud, in high-dimensional phase-space, that seeks to be robust to factors such as scale and resolution. Briefly, Mapper represents the underlying distribution of data in a metric space as an undirected graph, where each vertex comprises a non-exclusive subset of data points spanning a patch of phase space. An edge is drawn between each two vertices that share at least one data point (Fig. 1A), representing connectivity between patches. We complement Mapper with a novel graph-theoretical analysis using k-nearest neighbor (kNN) distance to estimate the density of data points over each patch of phase space represented by a vertex, determine local maxima, and define metastable community states (Fig. 1B). In contrast to established methods such as hierarchical clustering, our method preserves the notion of a continuous underlying density distribution, with the states representing a discrete coarse-graining, and recognizes low-density regions of phase space unassociated with any metastable state. In addition, it is possible for a data point to be associated with more than one vertex in the Mapper graph and thus with more than one state, allowing identification of samples that fall between or are in transition between metastable states.

**Figure 1:**
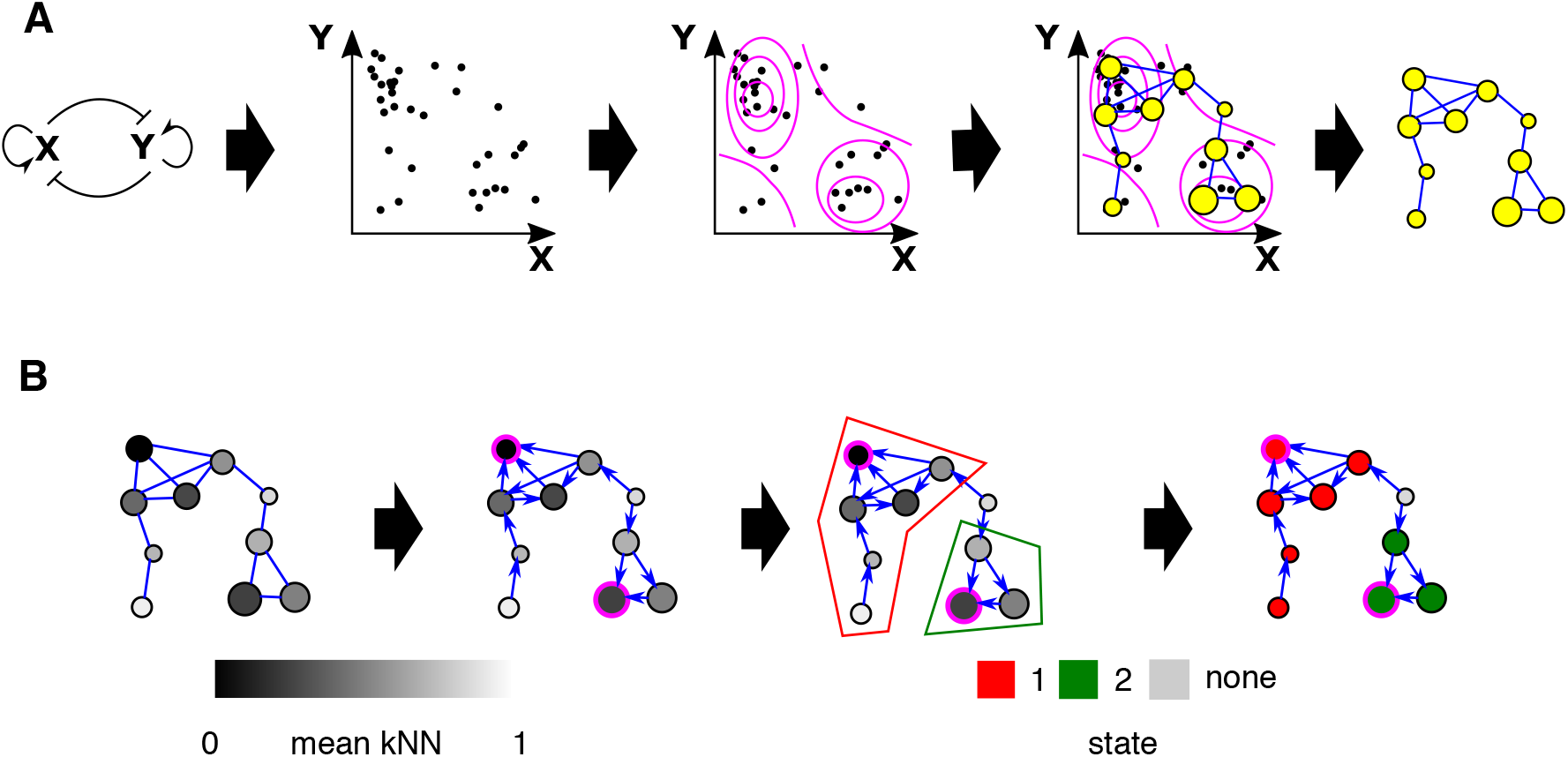
Using Mapper to characterize the microbial phase space. A. Cartoon of use of the Mapper algorithm to infer the probability density of a toy ecosystem. The mutually antagonistic interaction between species X and Y leads to denser sampling of the phase space where either X or Y is abundant and the other is rare than in other regions; configurations in which X and Y are similar in abundance are unstable, as small uncertainties in numerical advantage will eventually lead to the dominance of one species over the other. Mapper infers a ‘skeleton’ of density from the data represented as a point cloud. This representation preserves major features of the density such as the two densely-sampled clusters separated by a sparsely-sampled region. Size of vertices indicates number of data points aggregated in each vertex. B. Identification of local maxima and metastable states in the Mapper graph shown in A. Data density for each vertex is estimated by the inverse of the mean kNN distance (see Methods) for samples associated with that vertex. Shading indicates mean kNN distance over all data points included in a vertex. The graph is converted to a directed graph, with each edge pointing in the direction of increasing estimated density. A local maximum, highlighted in pink, is defined as a vertex that has higher density than all its neighbors. Finally, the state associated with a local maximum is defined as the set of vertices that have uniquely shortest directed graph distance to that maximum. Non-maxima vertices with equal graph distances to multiple local maxima are unassociated with any state (grey).

We used our method to infer the density and associated topological features of the point clouds for three published microbial time series data sets, two human gut microbiomes—one of stool samples collected from seven cholera patients from disease through recovery [23], one from two mostly healthy adult males [8]—and one of marine *Prochlorococcus* communities spanning multiple depths collected from one site in the Atlantic Ocean (BATS) and one in the Pacific (HOT) [32]. (For details on the sampling frequency and duration for each data set analyzed, see Supporting Information Table 1.) We selected these data sets in part to test our method by recapitulating biology known from the original studies, and in part to discover novel features not addressed by prior methods. In both human gut and marine systems, we find that significant physiological and environmental events, including recovery from infection and geochemical cycling, correspond to recurrent successions of state transitions. We show that these successions are an informative coarse-grained view of microbiome dynamics, with implications for the assessment of ecological resilience.

**Table 1:**
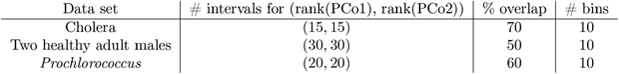
Hyperparameters used to generate the Mapper representation of each data set.

| Data set | # intervals for (rank(PCo1), rank(PCo2)) | % overlap | # bins |
| --- | --- | --- | --- |
| Cholera | (15, 15) | 70 | 10 |
| Two healthy adult males | (30, 30) | 50 | 10 |
| *Prochlorococcus* | (20, 20) | 60 | 10 |

## Results

### Dynamics of human gut microbiome recovery from cholera infection

We found the cholera phase space to be partitioned by clinical phenotype, i.e. diarrhea or recovery (Fig. 2A). Division of the phase space into states found that vertices within a state tended to consist of either samples from the diarrhea phase or from the recovery phase, rather than a mixture of both (Supporting Information Fig. 6). The original study [23] recognized phases of progression according to equal-time divisions of the diarrhea and recovery periods, respectively, of each patient. Our identification of disease substates,in contrast, is based on community composition and integrated across data from all patients. We found the diarrhea region was further subdivided into two states, 2 and 7 (Fig. 2B). Patients C, E, and G occupied state 7 for prolonged durations immediately before clinical recovery; patients A, B, and F stably occupied state 7 for approximately 20 hours, but switched to other states for the last few time points before clinical recovery (Fig. 2C). In the case of patient A, the final few time points were associated with state 5, which represented an intermediate region of the phase space between the diarrhea- and recovery-associated neighborhoods. These results suggest that state 2 constituted a universal ‘early’ diarrhea state, and state 7 a universal ‘late’ diarrhea state, with distinct community compositions. The original study noted taxa which consistently changed in abundance between the start and end of the diarrhea phase, for example *Streptococcus* and *Fusobacterium* [23], here we show that these compositional shifts are observable on the whole-community scale.

**Figure 2:**
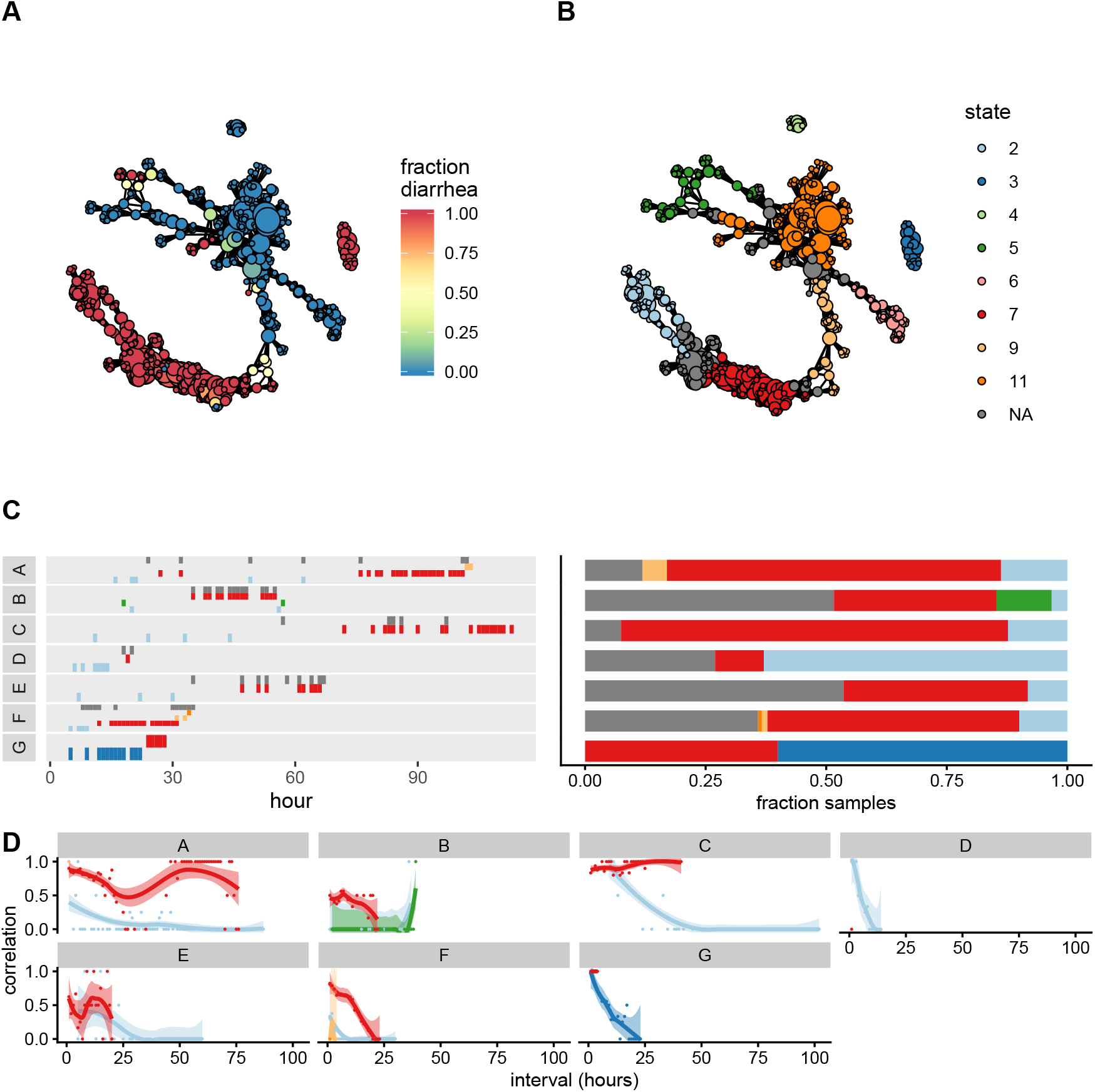
The phase space of the cholera gut microbiome. A. Mapper representation of the combined cholera data reveals disease- and healthy-associated neighborhoods of the phase space. Color: fraction of samples in each vertex associated with diarrhea. Connected components of the Mapper graph representing only one sample are not shown. Disjoint regions of phase space are represented as separate connected components. B. Partitioning of the phase space into metastable states. Vertices unassigned to any state are colored in grey. C. Left: progression of subject compositions during the diarrhea phase by state, showing persistence of states over time. Y axis and color indicate state index, with color indexing as in B. Where a sample was associated with multiple states, all were included. Right: frequency of samples associated with each states during the diarrhea phase for each subject with colors as in B. D. Temporal correlation function for the diarrhea phase of each subject. Dots: raw values of 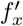 for pairs of samples (see Methods). Lines: smoothed empirical mean of 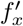. Ribbons: standard error of the mean. Values outside the range of 0 ≤ *y* ≤1 omitted.

Generally, patients occupied state 7 for longer than they did state 2, suggesting that the stability of the late state in a given patient influences disease duration. To quantify stability, we calculated a temporal correlation function for each state-patient pair during the diarrhea phase (see Methods). Monotonically decreasing correlation functions indicate metastability, showing that the system transiently occupies a state before transitioning to a different state; slopes become more negative with decreasing stability. While this analysis revealed that all patients transiently occupied state 2, with greatest persistence in patient C, patients A, C, and E had non-monotonic correlation functions for state 7, coinciding with prolonged times to recovery compared to the rest of the cohort, with patients B and F exhibiting the expected monotonic decrease (Fig. 2D). This indicated that patients A, C, and E repeatedly entered and exited state 7, suggesting that prolonged diarrhea in these three patients may have been additionally influenced by the instability or inaccessibility of alternative, healthy states, and that (re-)assembly of the healthy microbial community constitutes a non-trivial step in recovery.

### Dynamics of two healthy adult microbiomes with transient diarrhea

In contrast to the cholera data set, the two healthy adult gut microbiome time series from David *et al*. [8] were separated by subject (Fig. 3A). Despite being clinically healthy for most of the observation period, both subjects’ microbiomes experienced perturbations: sub ject A traveled from his residence in the United States to southeast Asia, twice experiencing traveller’s diarrhea; and subject B, also based in the US, suffered an acute infection by *Salmonella*. Previous studies [8, 19] noted that, while the microbiome of A returned to its original state after travel, recovery from *Salmonella* left the microbiome of B in an alternative state. Confirming this, we found that subject A occupied the same regions of phase space before and after travel, while subject B occupied disjoint regions before and after infection. We further found that the post-*Salmonella* samples of subject B distributed over several connected components, showing that the gut microbiome of subject B remained in flux across several distinct compositional substates even after being clinically marked as having recovered (Fig 3B). Division of the phase space into states found that vertices within a state tended to be dominated by samples from a single subject (Supporting Information Fig. 7).

**Figure 3:**
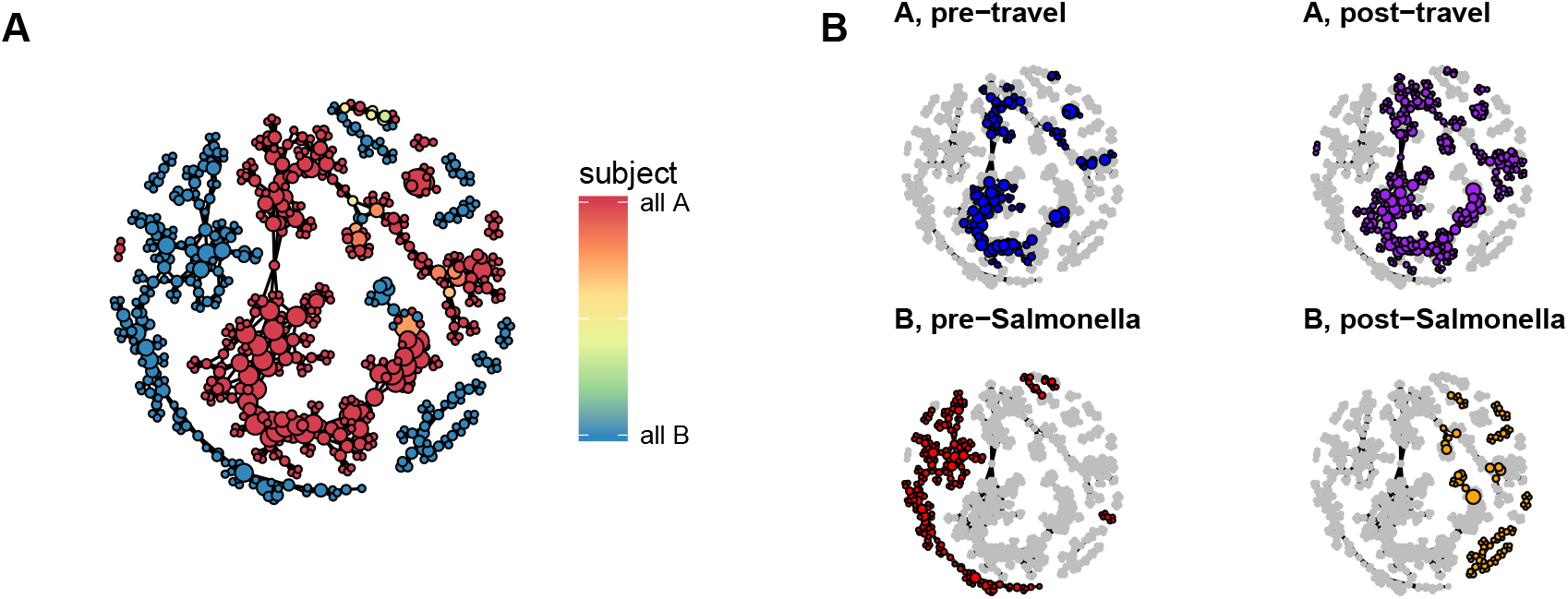
The phase space of two healthy adult male gut microbiomes. A. Mapper representation of the combined daily time series of two healthy adult human gut microbiomes. Connected components of the Mapper graph representing only one sample are not shown. B. Regions of phase space occupied by each subject before after perturbation.

The large connected components representing the pre- and post-travel healthy samples of subject A and the pre-Salmonella healthy samples of subject B were each divided into several states (Supporting Information Fig. 1), suggesting that the clinical ‘healthy’ phenotype of an individual is a probability over multiple compositionally distinct states. The existence of states in microbiome phase space proposes a novel metric for microbiome resilience: comparing the distribution of samples across states between time windows. Subject A occupied states with identical probability before and after travel, exhibiting resilience; in contrast, subject B post-infection did not restore the pre-infection probability across states, despite some samples sharing states with pre-infection healthy samples (Fig. 4A). Thus, the restoration of the microbial community to a ‘healthy’ state cannot be confirmed with a single time point.

**Figure 4:**
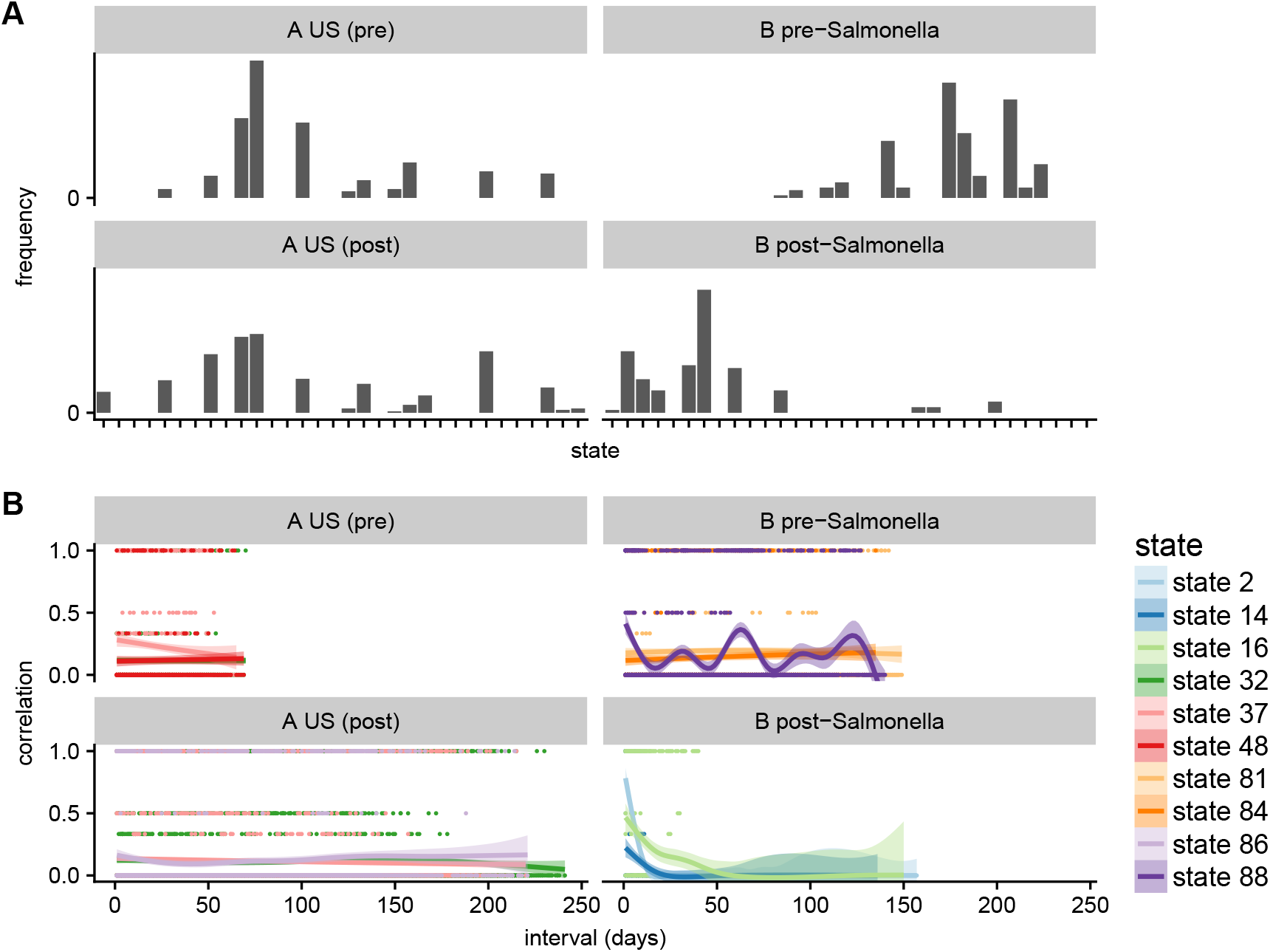
States and dynamics of two healthy adult male gut microbiomes. A. Frequency of states for healthy periods before and after perturbation. X axis: state index. Y axis: frequency of samples. B. Temporal correlation functions for the three most probable states during each event in the ‘healthy’ phases of each subject. Dots: raw values of 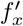 for pairs of samples. Lines: smoothed empirical mean of 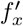. Ribbons: standard error of the mean.

Temporal correlation functions further showed that subject A, as well as subject B before infection, repeatedly visited the same set of states; in contrast, subject B after infection transiently occupied several states without repetition (Fig. 4B). This shows that not only did the microbiome of subject B enter an alternative state, or probability across states, post-infection, but that this alternative state was not fully stabilized. It is possible that the pre-infection probability across states was restored in subject B after the end of the observational period.

### Recurrent seasonal dynamics of *Prochlorococcus* communities in the Pacific and Atlantic

Compared to the phase spaces of human gut microbiomes, which may be discretized by individual or phenotype, the *Prochlorococcus* phase space was organized by gradients of depth (Fig. 5A) and temperature (Supporting Information Fig. 4), indicating that, in these environments, small changes to environmental conditions result in small changes to community structure. In contrast to the two human gut microbiome data sets, division of the *Prochlorococcus* phase space into states found the mean depth per vertex in each state to vary continuously (Supporting Information Fig. 8). The phase space possessed multiple states (Fig. 5B), with state 4 largely representing shallow fractions of the water column ≤ 100m; states 2, 3, and 6 deeper fractions; and state 1 intermediate depths. State 5 represented an infrequently-occupied region sampled only by the 140m fraction at BATS on January 27, 2004, and by the 125m fraction at HOT on January 31, 2008 (Fig. 5C). As such, state 5 possibly constitutes an alternative state for deep water fractions in mid-winter. Communities differing in depth rarely shared compositions, and transitioned between states, in many cases periodically across calendar years (Fig. 5C), showing that some communities experienced abrupt periodic shifts in environmental conditions due to geochemical events.

**Figure 5:**
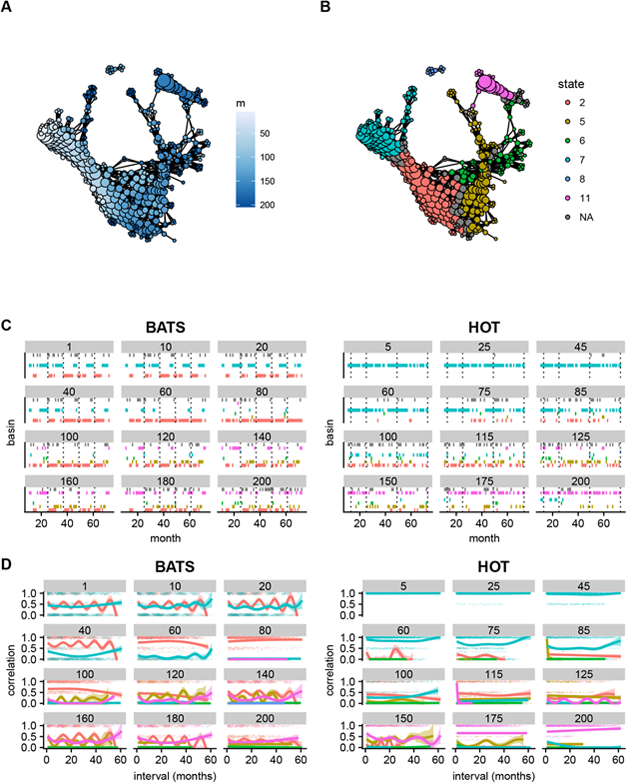
The combined phase space of two *Prochlorococcus* communities inhabiting the Atlantic and Pacific Oceans, respectively. Connected components of the Mapper graph representing only one sample are not shown. A. Vertices colored by mean depth in meters of represented samples. B. Partitioning of the phase space into states. C. Successions of states for each site-depth fraction combination. Dotted lines indicate samples during January. Colors indicate states as in B. D. Temporal correlation functions for each state per site-depth fraction combination. Dots: raw values of *f_x_* for pairs of samples. Lines: smoothed empirical mean of 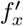. Ribbons: standard error of the mean.

Despite the graduated variation of composition with depth and temperature, the range of compositional dissimilarity across the range of environmental conditions is sufficient to constrain given depth fractions to a neighborhood of phase space, such that shallow- and deep-fraction *Prochlorococcus* communities rarely occupy the same compositional states over time (Fig. 5C). However, it is known that the BATS water column undergoes an annual late winter upwelling [32], intermixing communities that otherwise inhabit different depths, and homogenizing environmental conditions across depths. We predicted that mixing would drive communities at all depths at BATS to converge on a common state, while no convergence would be observed at HOT. Accordingly, we observed a transition to state 1 by all depths at BATS in January of each year. After June, depths 1-20m and 120-200m relax toward states characteristic of shallow and deep depth fractions, respectively, while state 1 persists longer in intermediate depths 40-100m. By contrast, no such upwelling occurs at HOT, and the probability of a given depth fraction occupying any state remains uniform over the calendar year; the distribution is especially stationary for shallow depths (Fig. 5C). This periodicity was also evident in periodic correlation functions for BATS, and non-periodic for HOT (Fig. 5D).

### Robustness of phase space characterization

Given that the data sets analyzed here are among the largest longitudinal microbiome data sets currently available, we asked whether the biological hypotheses could have been obtained from sparser data sets. We focused on our finding that microbiome phase spaces are structured by latent variables representing host phenotypes or environmental conditions, and examined whether this structuring was robust to data rarefaction. We found that the partitioning of the phase space by clinical phenotype in the case of the cholera patients, by subject in the case of the two healthy adult humans, and the gradation by depth in the case of *Prochlorococcus* communities, are robust to all rarefaction tests performed. In the case of cholera patients, nodes remained divided into those representing mostly samples from the diarrhea phase and those representing the recovery phase, with edges being more dense between nodes of the same phenotype than those of different phenotypes (Supporting Information Fig. 3). In the case of the two healthy adult humans, nodes were consistently dominated by samples from one subject, with edges being more dense between nodes representing the same subject than those representing different subjects (Supporting Information Fig. 4). For the *Prochlorococcus* data set, nodes aggregating samples from similar depth fractions were more densely connected than those representing disparate depths (Supporting Information Fig. 5).

### Comparison with hierarchical clustering and principal component analysis

To compare our method with standard methodologies, we performed hierarchical clustering and principal component analysis (PCA) on the OTU tables for each data set. We found that, while PCA confirmed the global partitioning of data by diarrhea or recovery within the cholera data set, partitioning by subject within the two adult gut microbiomes data set, and gradation by depth within the *Prochlorococcus* data set, it failed to make evident finer-grained features such as the existence of early- and late-diarrhea states in the cholera data set, and the distinction of pre- and post-*Salmonella* states for subject B in the two human gut microbiomes data set. Furthermore, the reduction of dimensionality to two dimensions through PCA made the visual separation between hierarchical clusters unclear, particularly for the cholera and two human gut microbiome data sets, and for the *Prochlorococcus* data set introduced a strong ‘horseshoe’ effect [33] (Supporting Information Fig. 9).

## Discussion

We identified unrecognized dynamics governing large-scale phenotypes in microbial time series data by using TDA to infer the shape of data density from 16S and ITS ribosomal RNA time series data. While analyses from the original studies identified bacterial taxa that were differed in abundance across host phenotypic or environmental states—for example the loss of *Firmicutes* in subject B post-*Salmonella* infection [8]—our method, by contrast, aims to identify transitions between global compositional states defined across all taxa without reference to metadata. Our results reveal the role of latent physiological and environmental variables [34], such as disease phenotype and phase of geochemical cycles, in organizing microbiomes over time. We observed common dynamics across instances of ecological processes in the two gut and one environmental timeseries datasets we studied. Using our approach, one can thus begin to infer general mechanisms that determine large scale phenotypes of clinical and environmental importance. The elements of our method— the definition of a metric phase space using the square root of the Jensen-Shannon divergence, the representation of the phase space using TDA, and the characterization of topological features using the adapted kNN density estimator and shortest graph distance searches—are specifically advantageous for analyzing high-dimensional compositional data. Relative abundances provide incomplete information on a system, and a system may be compositionally stable while remaining dynamic in absolute abundance [49]. Our method can be readily adapted to work with absolute abundance where such data are available. Compared to representational methods such as PCA, our method benefits from using all distance information; and compared to clustering techniques, our method does not require specifying the number of states, such as required in k-means.

While subjects in both human gut data sets experienced transient infection by bacterial pathogens, the large-scale dynamics differed between the two groups. We found that multiple cholera patients followed a trajectory of early-to late-stage disease states. In contrast, the two healthy subjects from the year-long data set experienced apparently random jumps between states during *Salmonella* infection and traveler’s diarrhea, respectively, that did not result in the stabilization in a reproducible alternate state during the course of disease. This discordance between the two human gut microbiome datasets suggests that microbial infections can potentially be classified into ‘ordered’ and ‘disordered’ types. Ordered infections are characterized by a reproducible trajectory through phase space, while disordered infections are characterized by unpredictable progression through phase space. The latter case represents a version of the ‘Anna Karenina principle,’ meaning individual microbiomes are more dissimilar during a particular perturbation than during health [51], while the former represents an inversion of the principle. Scale is likely important in this distinction: independent of the deterministic or stochastic nature of the perturbation induced by an infection, if its magnitude is smaller than ‘baseline’ fluctuations of the healthy microbiome, variations between individuals will remain the dominant variable in organizing the phase space. If the magnitude of the perturbation is larger, it may overwhelm individual variability and cause the phase space to instead appear organized by phenotype. Thus, data on the variability of healthy microbiomes over time between and within individuals will be crucial to characterizing the impact of a given disease on the microbiome. We also note that our conclusions are influenced by sampling frequency: our method cannot capture dynamics on a shorter time scale than that of sampling, and systems that seem noisy on a particular time scale may have ordered dynamics on longer time scales.

Our analysis of the David *et al*. data set shows that the microbiome of a healthy individual transitions between states over time. While key dominant taxa may persist, no single large-scale compositional state defines healthy physiology. However, an individual microbiome may occupy states with the same probability during two separate ‘healthy’ time windows. Integrating the information over time for each of the healthy periods, the physiological phenotype can be inferred to be stable despite the system state being dynamic. Put differently, if one interprets states as microstates of the microbiome composition, a systemic clinical or environmental phenotype could then be regarded as a macrostate, and a resilient ‘healthy’ microbiome will remain in a stable *macrostate* over time.

This notion of resilience as identical probability across states before and after a perturbation can be generalized to a notion of dynamic stability, defined as stationary probability across states over time. Dynamically stable microbiomes do not necessarily stabilize within a single state, but revisit a given set of states with fixed probability. Our temporal correlation analysis shows that dynamically stable microbiomes, such as subject A and subject B pre-infection from the study in [8], are characterized by non-monotonic temporal correlation functions, indicating the microbiome revisits the same states over time. In contrast, unstable microbiomes, such as subject B post-infection, exhibit monotonically decaying correlation functions, indicating the microbiome transiently occupies compositional states without recurrence. Dynamical instability can persist after infection even in the microbiome of an individual clinically marked as having recovered from infection, as in the case of subject B, revealing additional nuances to the association between stability and health in human microbiomes. The ability to assess resilience from data in the absence of detailed knowledge of the underlying network of microbe-microbe interactions complements model-based methods that analytically solve for fixed points and linear stability [5]. Alternate means of estimating stability and resilience may be possible, for example by quantifying the degree to which consecutive time points are associated with the same or adjacent Mapper vertices.

For the two human gut microbiome data sets, we observe some of the same phenomena as the original studies: for the seven cholera patients, certain taxa were differentially abundant throughout the progression of disease [23]; and for subject B of the two healthy males, that the pre-*Salmonella* microbiome composition was not recovered by the end of the experiment [8]. In the first case, we remark that differential abundance of individual taxa does not necessarily imply the existence of large-scale compositional states consistent across patients and disease phases, such as we describe here. In the second case, we additionally found multiple states in the pre- and post-perturbation healthy phases of both subjects, and showed that restoration of a healthy and resilient microbiome is associated with the recovery not of a specific composition but of a distribution across compositional states.

We point out several caveats regarding our method. First, though we defined the phase space using the Jensen-Shannon distance, other metrics may be used, and the results of analysis using different metrics for the same data should be compared in future applications. Second, due to the lack of an established protocol for selecting Mapper hyperparameters, we used a heuristic method to choose their values for our analyses. A more rigorous optimization method is desirable, especially one developed against synthetic data from *de novo* simulations where the ‘ground truth’ of the parameters, and thus the shape of the density, are known *a priori*. Third, we use Mapper to create a representation of the density, but question of whether it is effective to analyze microbiome dynamics via the topology of the density in a given case is independent of Mapper and TDA, and other methods may be used. Fourth, we assume the data accurately represent the compositions of the sampled communities, when in fact challenges exist with translating sequencing data into compositions [18, 17]; addressing these challenges is outside the scope of this manuscript.

In addition to offering a novel quantitative description of microbiome states and dynamics, we hope our analysis will, in time, facilitate predictive modeling of the dynamics and forecasting of major state transitions in the microbiome. As an example, our approach to identifying states from microbial time series can be used to infer state transition probabilities under different conditions, and thus can serve as a basis for fitting the parameters of Markov chain models [9, 12]. The concept of the potential landscape that motivated our study is closely linked to the theory of critical transition forecasting [6, 7, 29, 40, 42]: as perturbations destabilize a system, it ascends the potential gradient and eventually reaches a tipping point from where it can rapidly enter into an alternative stable state. Topological analyses, in turn, may eventually facilitate characterization of the potential landscape based on past observations, and real-time estimation of its stability and state transition probability. Both of these approaches allow modeling and prediction of major dynamical events without detailed knowledge of underlying mechanisms, and may prove pivotal to understanding complex, data-rich biological systems not limited to microbiomes, but also including, for instance, gene regulatory networks and animal ecosystems.

## Methods

### Human gut microbiome data and preprocessing

The publicly available data that we re-analyzed here were generated by David *et al* [8] accessible on the European Nucleotide Archive (ENA) under the accession number ERP006059, and by Hsiao *et al* [23] on the NCBI Short Read Archive (SRA) under the accession number PRJEB6358. The downloaded reads were trimmed with V-xtractor version 2.1 [21] (a HMM scan based method of isolating variable regions from 16S rRNA sequences) to ensure the amplicon sequences could be aligned across consistent fractions of the 16S rRNA variable regions. Trimmed reads were then clustered into OTUs using usearch v9.2.64 [11] with a minimum cluster size of two. Representative sequences from each OTU were classified using mothur v1.36.1 [43] and the RDP reference 16S rRNA sequences v16 [4].

### Prochlorococcus data

Data from Malstrom *et al* [32] was obtained from the Biological and Chemical Oceanography Data Management Office (https://www.bco-dmo.org), accession number 3381.

### Mapper

Conceptually, the Mapper algorithm accepts as input a matrix of distances or dissimilarities between data, and aims to represent the shape of the distribution of data points in high-dimensional phase space as an undirected graph. In this graph, vertices represent neighborhoods of phase space spanned by subsets of adjacent data points, and edges represent connectivity between neighborhoods. In brief, it does this by dividing the data into overlapping subsets that are similar according to the output of at least one filter function that assigns a scalar value to each data point, performing local clustering on each subset, and representing the result as an undirected graph, where each vertex represents a local cluster of data points, and edges between vertices represent at least one shared data point between clusters.

### Distance matrix

We interpreted microbiome relative abundances to be probability distributions, and thus used the square root of the Jensen-Shannon divergence as a metric [27]. However, it is important to note that any other metric can be used in place of the Jensen-Shannon distance, such as the Aitchison distance [37], calculated from centered [28] or isometric [44] log-transformed relative abundances.

### Filter functions and binning

For the filter functions used by Mapper to bin data points, we performed principal coordinate analysis (PCoA, also known as classical multidimensional scaling) in two dimensions on the pairwise distance matrix, and used the ranked values of principal coordinates (PCo) 1 and 2 as the first and second filter values for Mapper, following Rizvi *et al*. [39]. PCo ranks are an appropriate filter for our purposes, as it assigns similar filter values to points that are relatively close together in the original phase space. We wish to note that while PCoA leads to loss of information, the following local clustering step is performed using subsets of distances from the original distance matrix, and is thus not affected. The data points were then binned by overlapping intervals of the two ranked principal coordinates. For hyperparameters specifying these bins and their overlaps, see Table 1.

### Local clustering

The algorithm first performs hierarchical clustering from all pairwise distances between data points within a bin of filter values. Then, it creates a histogram of branch lengths using a predefined number of bins, and uses the first empty bin in the histogram as a cutoff value, separating the hierarchical tree into single-linkage clusters. The algorithm thus finds a separation of length scales within each neighborhood of phase space represented by a bin of the filter values. We used the default number of histogram bins, 10, for each data set (Table 1).

### Creating the undirected Mapper graph

The final output is produced by representing each local cluster of data points as a vertex, and drawing an edge between each pair of vertices that share at least one data point. When plotting, the size of each vertex represents the number of data points therein.

### Selection of hyperparameters

The Mapper algorithm is relatively new, and there are currently no standard protocols to optimize the values of the hyperparameters. For our purposes, it was important that the algorithm achieved a sufficiently high resolution in partitioning data, but also adequately represented connections between regions of phase space. We thus used the following heuristic to set the number of intervals and percent overlap for each data set.

1. The largest vertex in the resultant Mapper graph should represent no more than ≈ 10% of the total number of data points in the set;
2. the number of connected components representing only one data point should be minimized.

We acknowledge that a heuristic determination of appropriate hyperparameter values leaves much to be desired; as such, we recommend future in-depth theoretical explorations of how the Mapper output depends on the choice of hyperparameters.

### Density estimation

We estimated the inverse density for each vertex by calculating the *k*-nearest neighbors (kNN) distance [10] for each constituent data point *i*:

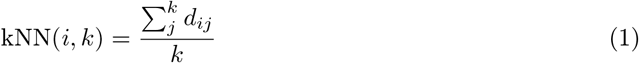

where *d_ij_* is the distance between points *i* and *j*, choosing *k* equal to 10% of the number of samples in each data set, rounded to the nearest integer. For a vertex *V* representing *n* points, we define its inverse density as

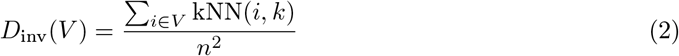

The *n*^2^ term in the denominator compensates for the differing sizes of vertices. Finally, we invert the inverse density to obtain the estimated density:

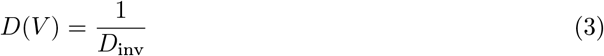

### State assignment

We then defined states as topological features of the density surrounding local maxima of *D*. We designated each vertex with higher *D* than its neighbors to be a local maximum of the potential. Connected vertices tied for maximum *D* were each assigned to be a local maximum. To approximate a gradient, we converted the undirected Mapper graph to a directed graph, with each edge pointing from the the vertex with lower *D* to the one with higher *D*. For each non-maximum vertex, we found the graph distance *d_g_* to each local maximum constrained by edge direction. We defined the state *B_x_* of a maximum *V_x_* as the set of vertices *V* with uniquely shortest graph distance to *V_x_*:

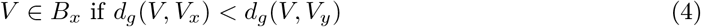

for all *y* ≠ *x* and *V_y_* ∈ *M*, where *M* is the set of all local maxima (Fig 1B). Vertices equidistant to multiple maxima were defined to be unstable regions unassigned to any state. Multiple connected maxima were defined as belonging to the same state. Notably, one data point may be associated with multiple vertices and states, or an unstable region and at least one state: we interpreted this to mean that the point is near a saddle point separating states, and as the ‘true’ coordinates of the saddle point are unknown, the data point is assigned to *all* such states and/or an unstable region with uniform weight.

### Calculating the temporal correlation function

Given that a system occupied state *B_x_* at time *t*, we defined the temporal correlation to be the probability that it will still (or again) occupy state *B_x_* at time *t* + *τ*:

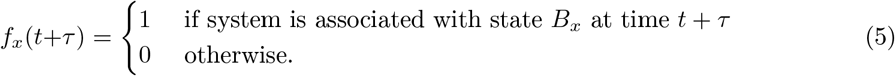

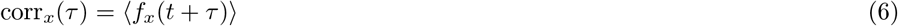

We calculated the correlation function for each state *x* visited by a subject during a characteristic period and for all sampled intervals between pairs of samples of length *τ*, where the subject was in state *B_x_* in the sample at the start of the interval. For the cholera data set, we calculated correlation functions for each state visited by each subject over the disease period. For the data set of two healthy adult males, we calculated correlation functions for each state visited by each subject in each healthy period, either before or after infection. For the *Prochlorococcus* data set, we calculated correlation functions for each state at each depth fraction at either site. Where a data point is associated with multiple states, we weigh the association with each state as 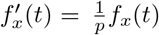, with *p* the total number of unique states associated with the system at time *t*, with the unassigned/unstable state regarded as a single distinct state. Notably, this means 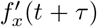 can have values of 1, 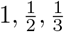…

### Rarefaction test

We created random subsets of each data set representing 90%, 50%, and 10% of the original data points, repeating 10 times for each data set and downsampling ratio. We then created Mapper graphs representing the rarefied data using the same hyperparameters as for each of the full data sets. We colored the vertices to indicate the same features as for the full data sets: for the cholera data set, by fraction of samples belonging to the diarrhea or recovery phase; for the two healthy adult gut microbiomes data set, by fraction of samples obtained from each subject; and for the *Prochlorococcus* data set, by the mean depth from which samples originated. We ordered the vertices by feature value and used a circularized linear layout algorithm, such that vertices with similar feature values are adjacent. Finally, we used shading to display edge densities.

### Software and data

The main repository for the study can be found on GitHub, at http://github.com/kellylab/microbial-landscapes.

An open-source implementation of Mapper in R, TDAmapper, was used for the main analysis and can be found at http://github.com/wkc1986/TDAmapper. This package was forked from the original implemented by Daniel Müllner which is maintained by Paul T. Pearson and can be found at https://github.com/paultpearson/TDAmapper.

## Supporting information

Cholera state mean taxonomy

Two human adult gut microbiome state mean taxonomy

Prochlorococcus state mean taxonomy

## Author’s contributions

W.K.C. designed and performed the analysis. D.V. processed and performed OTU calling on the data from Hsiao *et al*.[23] and David *et al*.[8]. W.K.C., D.V., and L.K. wrote the manuscript.

## Additional Files

### Supporting information

- Supporting methods describing PCA and hierarchical clustering.
- Supporting table showing sampling frequency and duration for each of the data sets analyzed.
- Supporting figure showing the states of the two human gut microbiomes data set.
- Supporting figure showing the temperature gradients across the *Prochlorococcus* phase space.
- Supporting figures showing the results of the data rarefaction test.
- Supporting figures showing the mean physiological or environmental properties per state for each data set.
- Supporting figure showing the results of PCA and hierarchical clustering.

### Supporting data

- Taxonomy tables showing the mean composition of each state for each data set.

