## Supporting information for "Topological analysis reveals state transitions in human gut and marine bacterial communities"

March 12, 2020

#### 1 Methods

##### 1.1 PCA and hierarchical clustering

PCA plots were generated using the prcomp function in the R package vegan v2.5-4 [1] and visualized using the R package ggplot2 v3.2.1 [2]. To perform heirarchical clustering analysis, bray-curtis dissimilarity values between samples within each dataset were calculated using the function vegdist in the R package vegan v2.5-4, and clustered using the R function hclust.

#### 2 Tables

| Data set | Sampling frequency | Duration | Notes |
|---|---|---|---|
| Cholera | irregular: multiple times/day during diarrhea, daily for first week after discharge, weekly over next 3 weeks of recovery, monthly over next 2 months | 33-76 hours diarrhea, 86-88 days recovery | 7 patients |
| Two adult human gut microbiomes | ∼ daily | 252 days (subject B)-364 days (subject A) | 2 subjects |
| *Prochloroccocus* | ∼ monthly | ∼ 5 years | 2 sites, 12 depth fractions each |

Table 1: Description of sampling frequencies and experiment duration for all data sets analyzed.

### 3 Figures

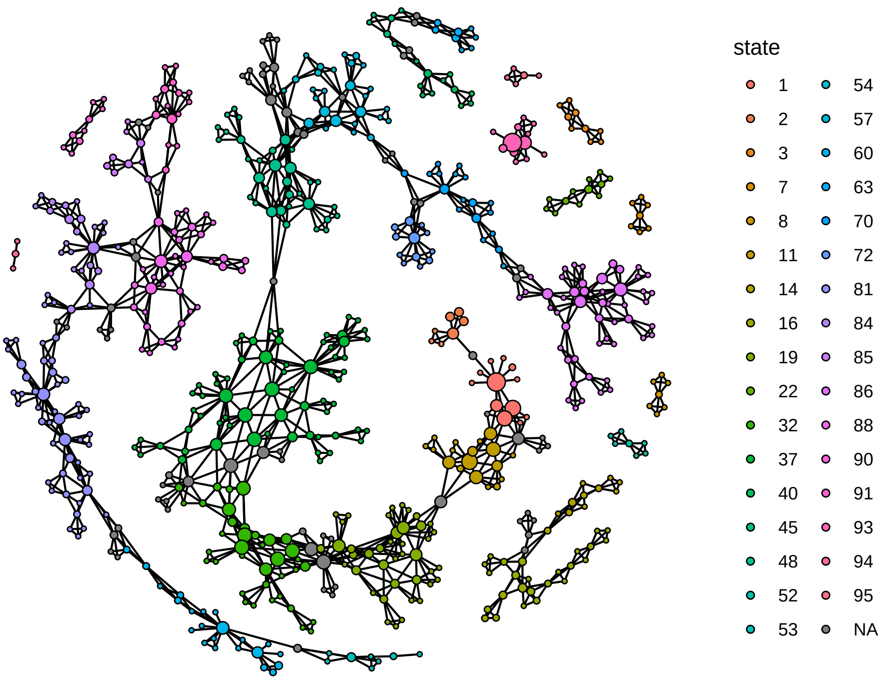


Figure 1: Metastable states for the data set of two healthy adult male gut microbiomes.

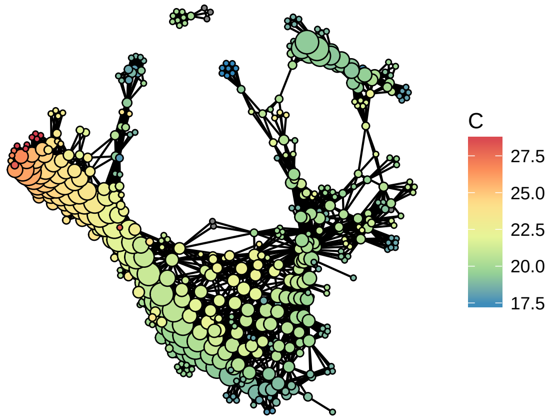


Figure 2: Mapper representation of the *Prochlorococcus* phase space colored by mean temperature per vertex. The phase space is spanned by a continuous temperature gradient.

.

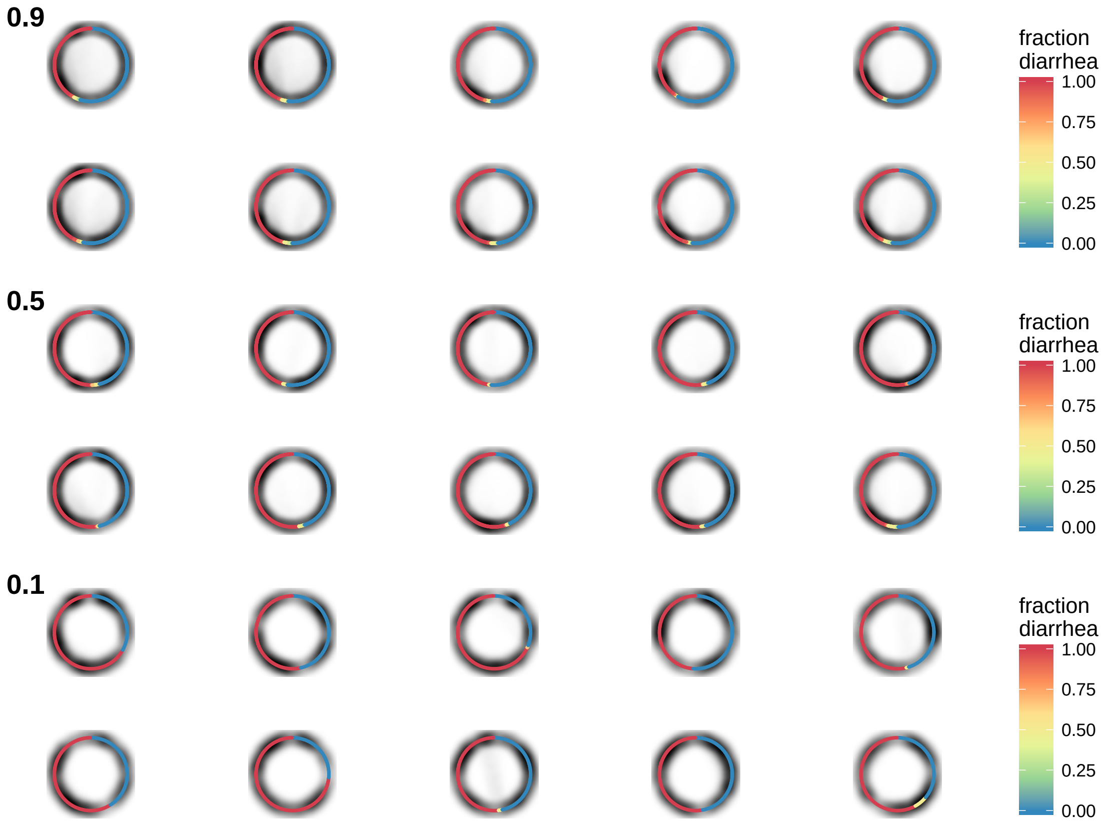


Figure 3: Mapper representations of 10 random subsets of the cholera data constituting 0.9 (top), 0.5 (middle), and 0.1 (bottom) of total data points, each. Nodes of Mapper graph laid out linearly and ordered by fraction of samples corresponding to phenotype, from 0% diarrhea (blue) to 100% (red). Gradient represents edge density. The majority of edges link nodes of similar phenotype, with few connections between the diarrhea and recovery regions of phase space.

.

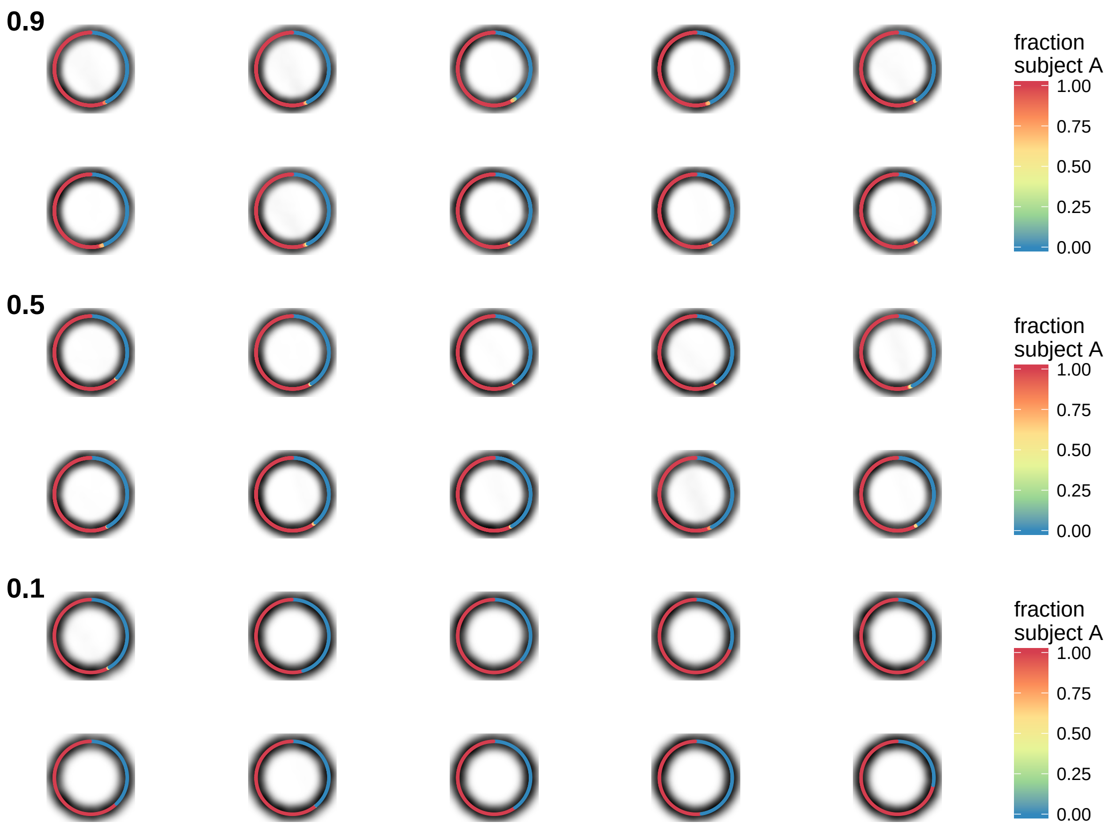


Figure 4: Mapper representations of 10 random subsets of the healthy human adult data constituting 0.9 (top), 0.5 (middle), and 0.1 (bottom) of total data points, each. Nodes of Mapper graph laid out linearly and ordered by fraction of samples corresponding to subject, from 0% subject A (blue) to 100% (red). Gradient represents edge density. The majority of edges link nodes representing the same subject.

.

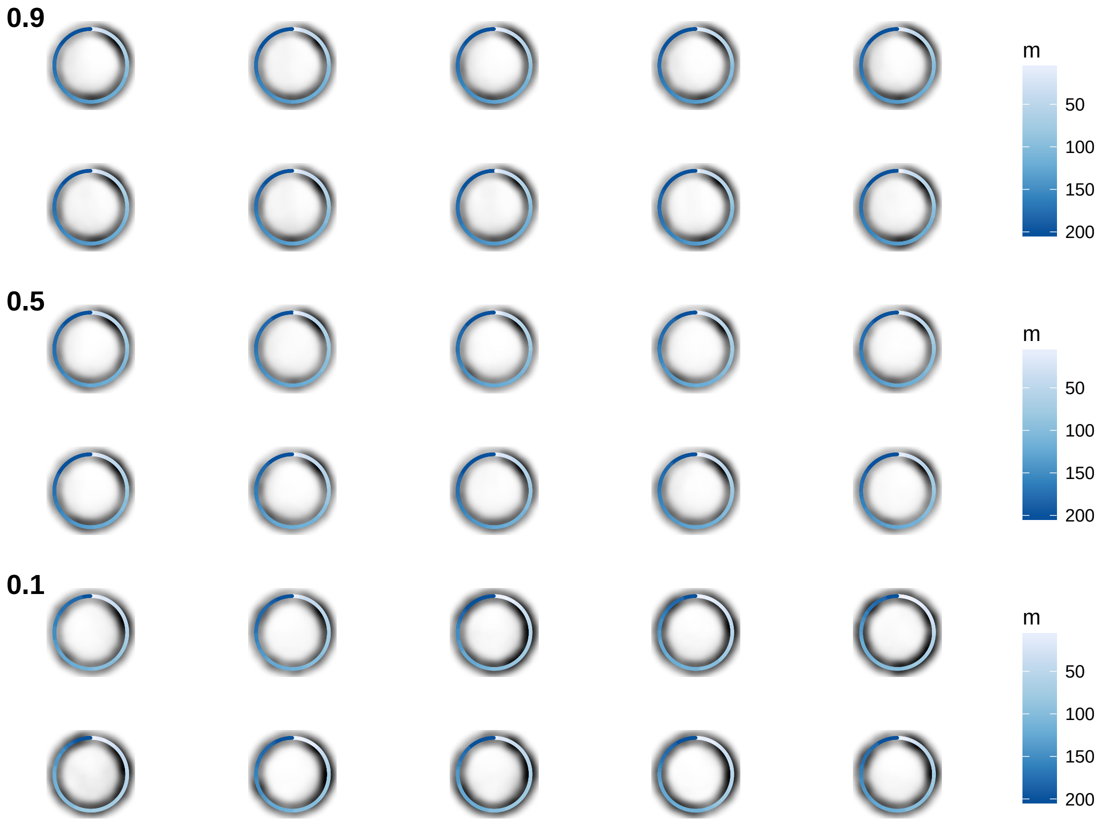


Figure 5: Mapper representations of 10 random subsets of the *Prochlorococcus* data constituting 0.9 (top), 0.5 (middle), and 0.1 (bottom) of total data points, each. Nodes of Mapper graph laid out linearly and ordered by mean depth, from shallow (white) to deep (blue). Gradient represents edge density. The majority of edges link nodes with similar depths,

.

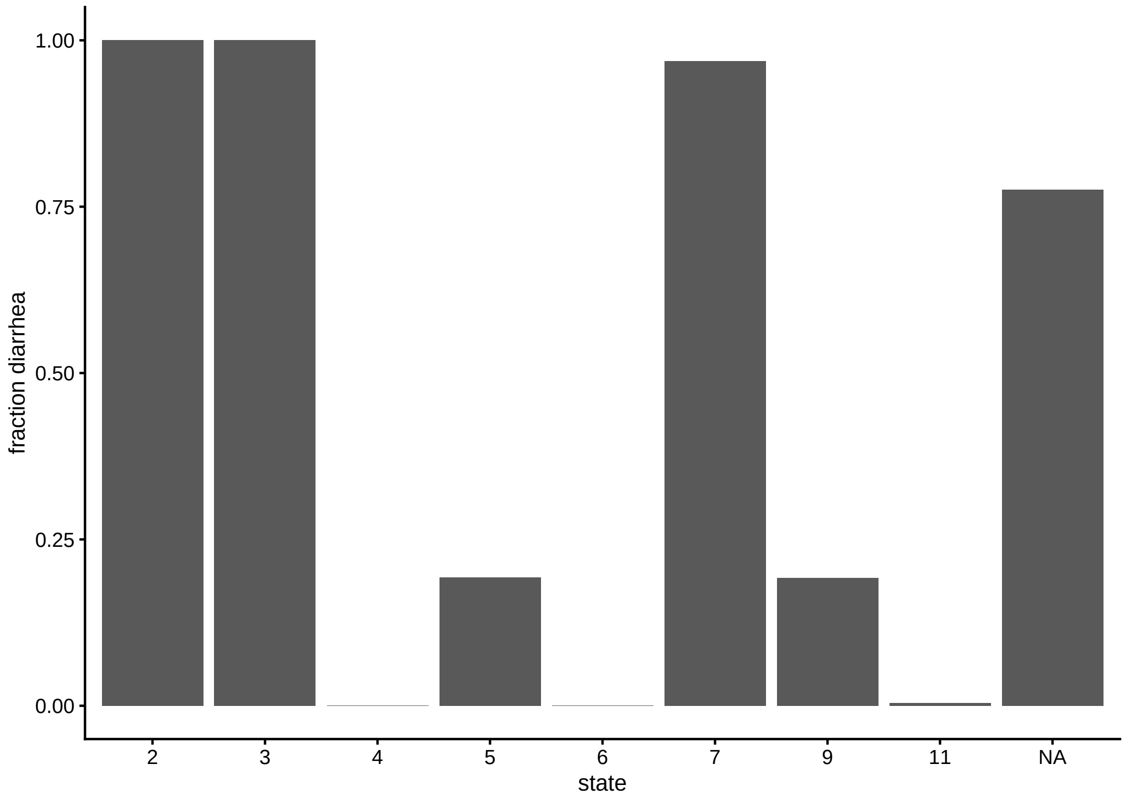


Figure 6: Mean fraction diarrhea per vertex for each state for the cholera data set.

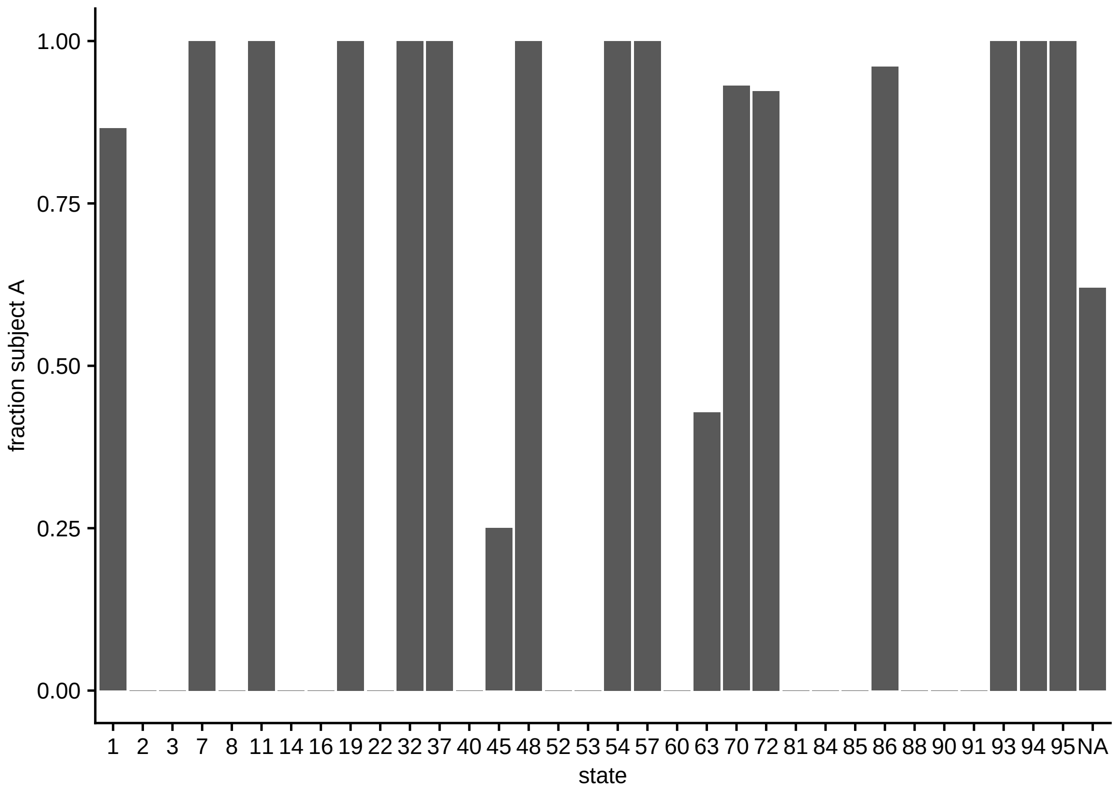


Figure 7: Mean fraction subject A per vertex for each state for the two adult gut microbiomes data set.

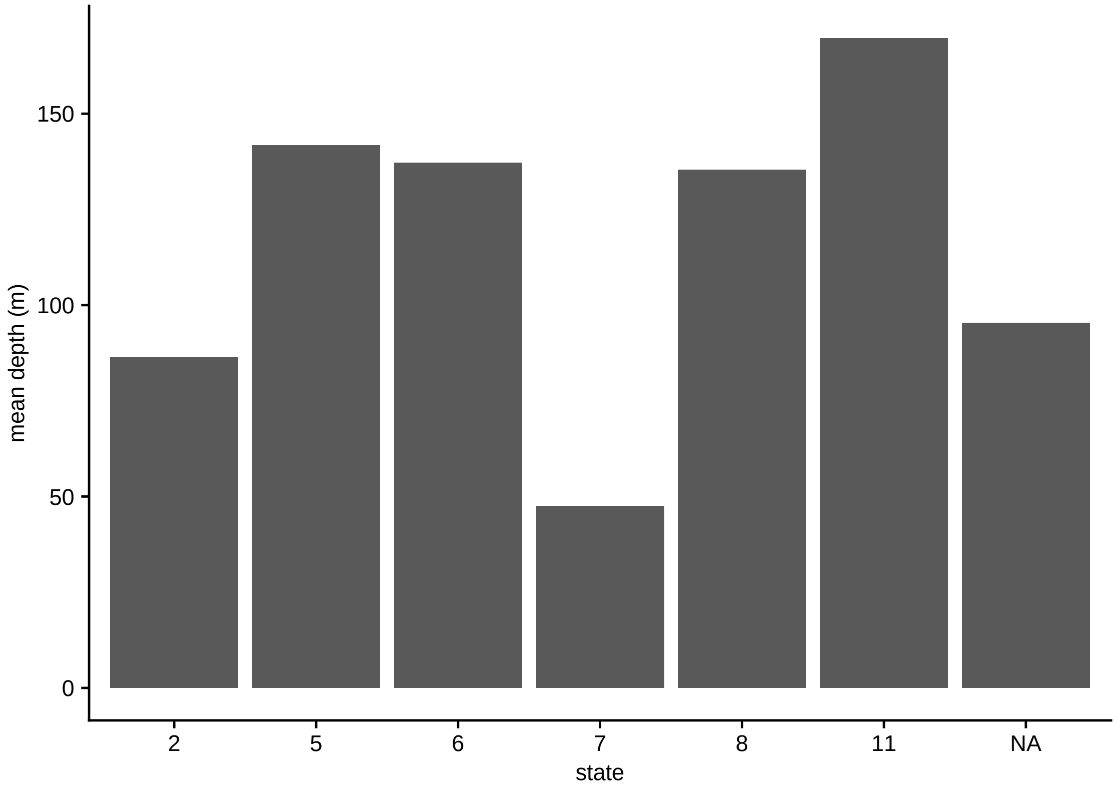


Figure 8: Mean depth per vertex for each state for the *Prochlorococcus* data set.

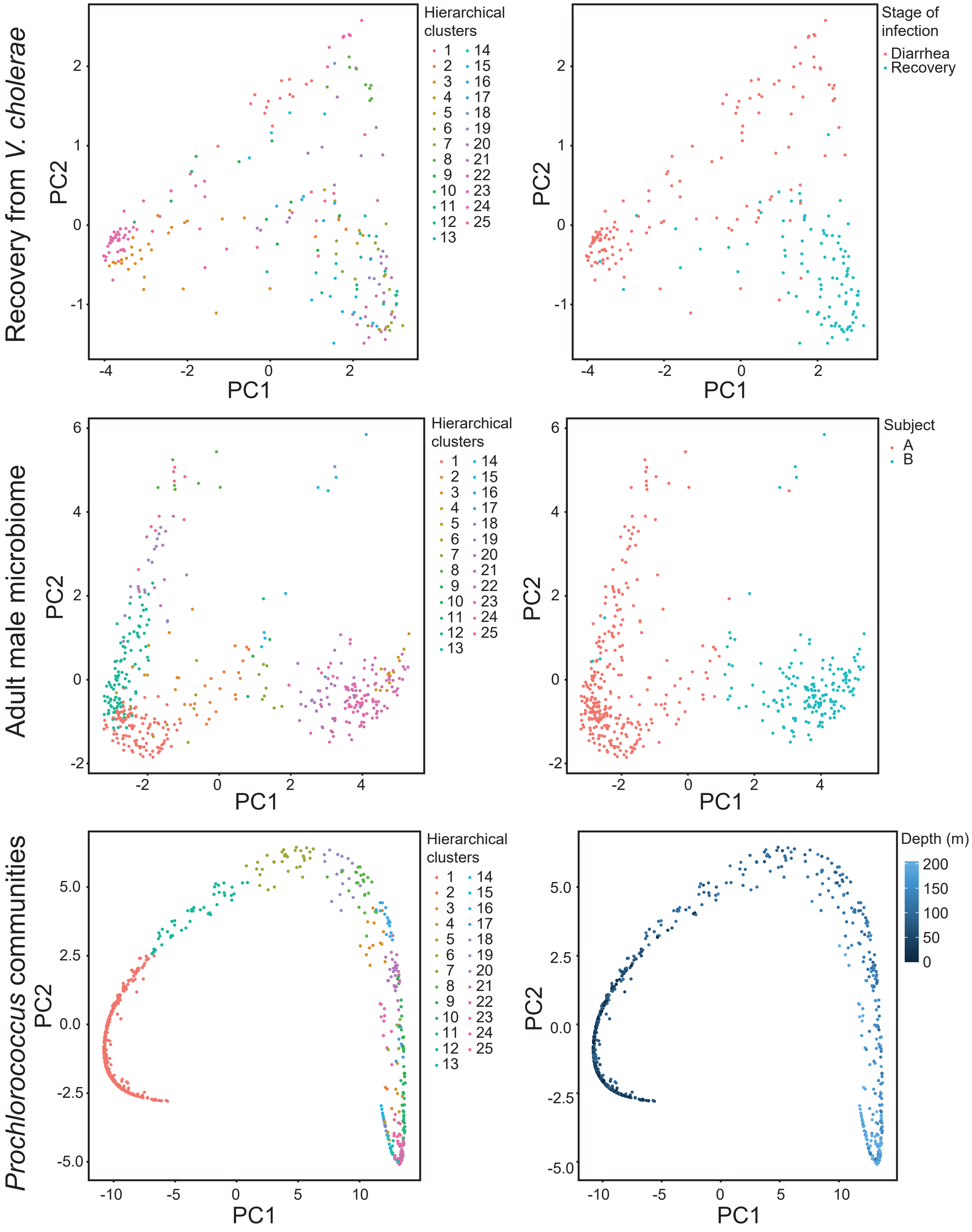


Figure 9: Principal component analysis of time series microbial communities. Samples are colored according to hierarchical clusters (left) and attributes of the person or depth they were taken from.
